# The Guaymas Basin Subseafloor Sedimentary Archaeome Reflects Complex Environmental Histories

**DOI:** 10.1101/2020.06.01.109066

**Authors:** Gustavo A. Ramírez, Luke J. McKay, Matthew W. Fields, Andrew Buckley, Carlos Mortera, Christian Hensen, Ana Christina Ravelo, Andreas P. Teske

## Abstract

We explore archaeal distribution and environmental niche differentiation in sedimentary subseafloor habitats of Guaymas Basin and the adjacent Sonora Margin, located in the Gulf of California, México. Specifically, we survey diverse subseafloor habitats on the Guaymas Basin flanking regions that are extending from the spreading center, termed here “off-axis” sites. Sampling locations include (i) control sediments without hydrothermal or seep influence, (ii) Sonora Margin sediments underlying oxygen minimum zone water, (iii) compacted, highly reduced sediments from a pressure ridge with numerous seeps at the base of the Sonora Margin, and (iv) sediments impacted by hydrothermal circulation at the off-axis Ringvent site. Generally, archaeal 16S rRNA gene datasets are largely comprised of Bathyarchaeal lineages, members of the Hadesarchaea, MBG-D, TMEG, and ANME-1 groups. The most frequently observed 25 OTUs belong to members of these lineages, and correspond to approx. 40 to 80% of the sequence dataset in each sediment sample. Differential distribution patterns of these archaeal groups in downcore sediments uniquely characterize each major sedimentary environment. Variations in archaeal community composition reflect locally specific environmental challenges throughout the greater Guaymas Basin area. Background sediments are divided into surface and subsurface niches, reflecting increased selection of the archaeal community downcore. In sum, the environmental setting and history of a particular site, not isolated biogeochemical properties out of context, control the subseafloor archaeal communities in Guaymas Basin and Sonora Margin sediments.

## Introduction

Guaymas Basin, located in the Gulf of California, México, is a young marginal rift basin where active seafloor spreading generates northeast-to-southwest trending axial troughs surrounded on both sides by extensive flanking regions (Lizarralde et al., 2007). In contrast to mid-ocean spreading centers, axial troughs and flanking regions of Guaymas Basin are covered by thick, organic-rich sediments that represent a combination of terrigenous input and biogenic sedimentation from the highly productive water column (Calvert, 1966). Magmatic intrusions, or sills, are embedded within the thick sediment layers, where they drive hydrothermal circulation (Lonsdale and Becker, 1985) and thermally alter buried organic matter (Seewald et al., 1990), in the process generating complex petroleum compounds (Didyk and Simoneit, 1989), light hydrocarbons and methane (Welhan and Lupton, 1987), carboxylic acids (Martens, 1990), and ammonia (Von Damm et al., 1985). Since the sediments act as a heat-retaining thermal blanket, magmatic activity and organic matter alteration and mobilization are not limited to the spreading center but also occur at considerable distance, up to 50 km off-axis (Lizarralde et al., 2010). Many of these off-axis sites resemble cold seeps, where methane advection is linked to pathways formed by deeply buried magmatic sills (Geilert et al., 2018). If the underlying sill is sufficiently shallow and hot, the hydrothermal underpinnings of these off-axis sites becomes visible; the recently described Ringvent site provides an example (Teske et al., 2019).

In contrast, the Sonora Margin harbors classical cold seeps where sediment compaction drives reducing, methane-rich seep fluids to the surface. Numerous seep sites with carbonate outcrops and cold seep fauna have been observed on an eroding pressure ridge that follows the transform fault at the base of the Sonora Margin (Simoneit et al., 1990;Paull et al., 2007); the seep communities at these sites are largely based on methanotrophy and sulfide oxidation (Portail et al., 2015). Seep communities and sulfide-oxidizing microbial mats are also widespread on the Sonora Margin slopes (Vigneron et al., 2014;Cruaud et al., 2017).

Finally, most of the extensive flanking regions of Guaymas Basin and the Sonora Margin slope are covered by sediments without particular seep or hydrothermal influence; these sediments consist of mixed terrigenous runoff and biogenic components, dominated by diatoms (Calvert, 1966). Sediments on the upper Sonora Margin underlying the oxygen minimum at ca. 600 to 800 m depth, lack bioturbation and show finely laminated, seasonally changing sedimentation patterns of spring diatom blooms and terrestrial runoff during late summer trains (Calvert, 1964).

Here, we survey the distribution of archaea in diverse sedimentary environments located in the greater Northern Guaymas Basin and Sonora Margin regions. Sampling areas include background sediments from the Guaymas Basin flanking regions, Sonora Margin sediment within the oxygen minimum zone, reducing sediment with cold seep characteristics from the base of the Sonora Margin, and sediment from the off-axis Ringvent site where hydrothermal circulation and methane seepage is driven by a gradually cooling, buried shallow sill. We expand a previous limited sequencing survey of these sediments focused on just one of these sites (Teske et al., 2019) by: i) extending the geochemical analyses, ii) increasing the sampling resolution used for molecular sequencing (from one or two samples per site to ~1-meter intervals for all sites), iii) providing a wide breadth of comparative ecological analyses, and iv) discussing the potential implications of our results at a basin-wide scale.

## Results

### 1. Sediment and porewater geochemistry

We surveyed archaeal distribution at six sites on the northwestern and southeastern off-axis regions of Guaymas Basin and on the Upper Sonora Margin (Figure 1, Table S1). These locations represent four different environmental settings: (i) Sediments on the Guaymas Basin flanking regions without hydrothermal or seep activity, represented by cores Cont03, Cont10 and Cont13; (ii) the oxygen minimum zone on the upper Sonora Margin (Calvert, 1964), represented by core OMZP12, (iii) compacted, highly reducing seep sediments from a pressure ridge, running along the transform fault that is cutting across the base of the Sonora Margin (Simoneit et al. 1990, Paull et al. 2007), represented by core Seep06, and (iv) the Ringvent site, characterized by off-axis hydrothermal circulation (Teske et al., 2019), represented by core RNVP11 (Figure 1). At each site, sediment piston cores ranging from 5 to 486 cm below the seafloor (cmbsf) were collected and geochemically characterized (Figure 2). The sediments are geologically young, ranging in age between ~0.05K and ~20K calendar years, as determined by C14 dating (Teske et al., 2019). The different cores show distinct geochemical characteristics.

**Figure 1:**
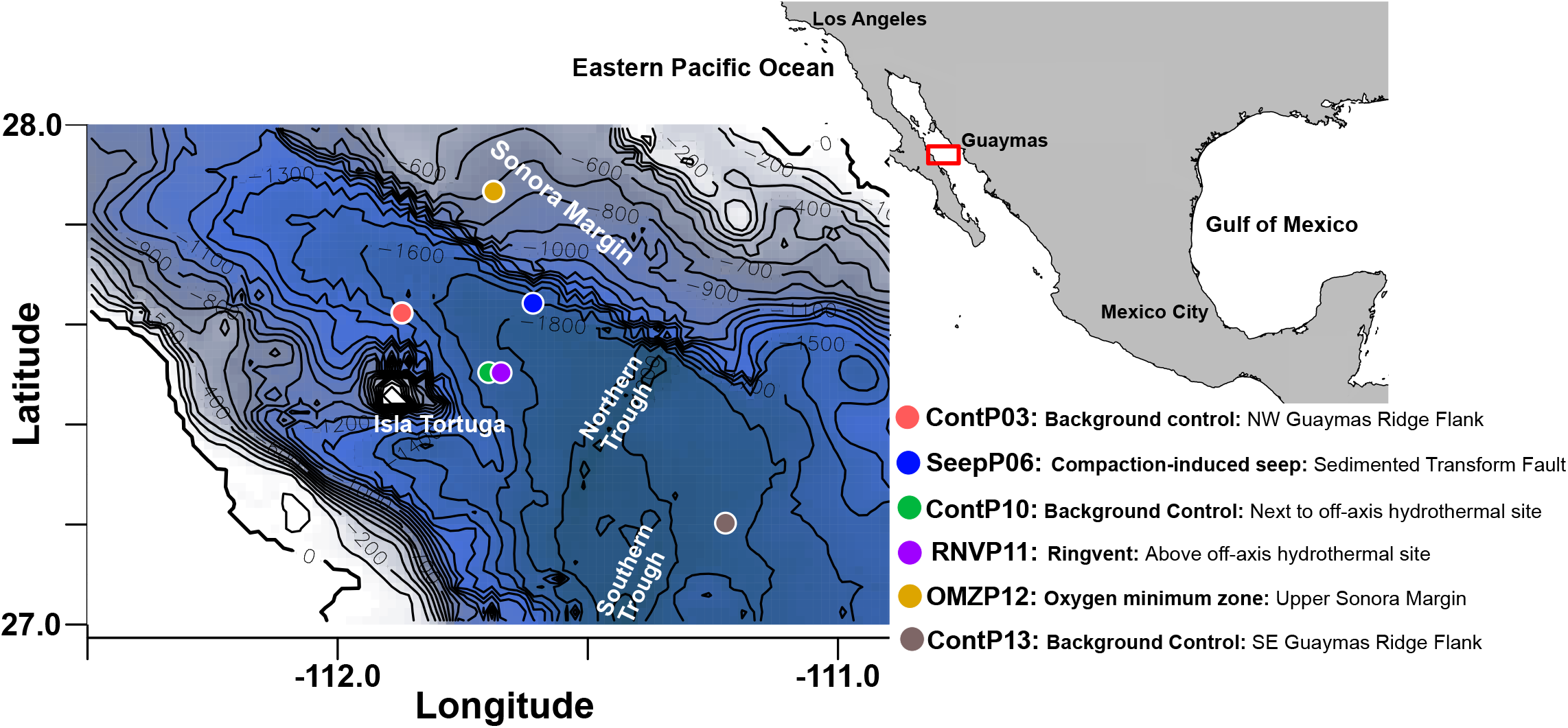
Continental and bathymetric hybrid map depicting the location of Guaymas Basin and Sonora Margin in the Gulf of California, and relevant coring sites of the *El Puma* cruise. The bathymetry blue scale is annotated with 100-meter isobaths; the deepest areas in the axial valley range to just below 2000 meters.

**Figure 2:**
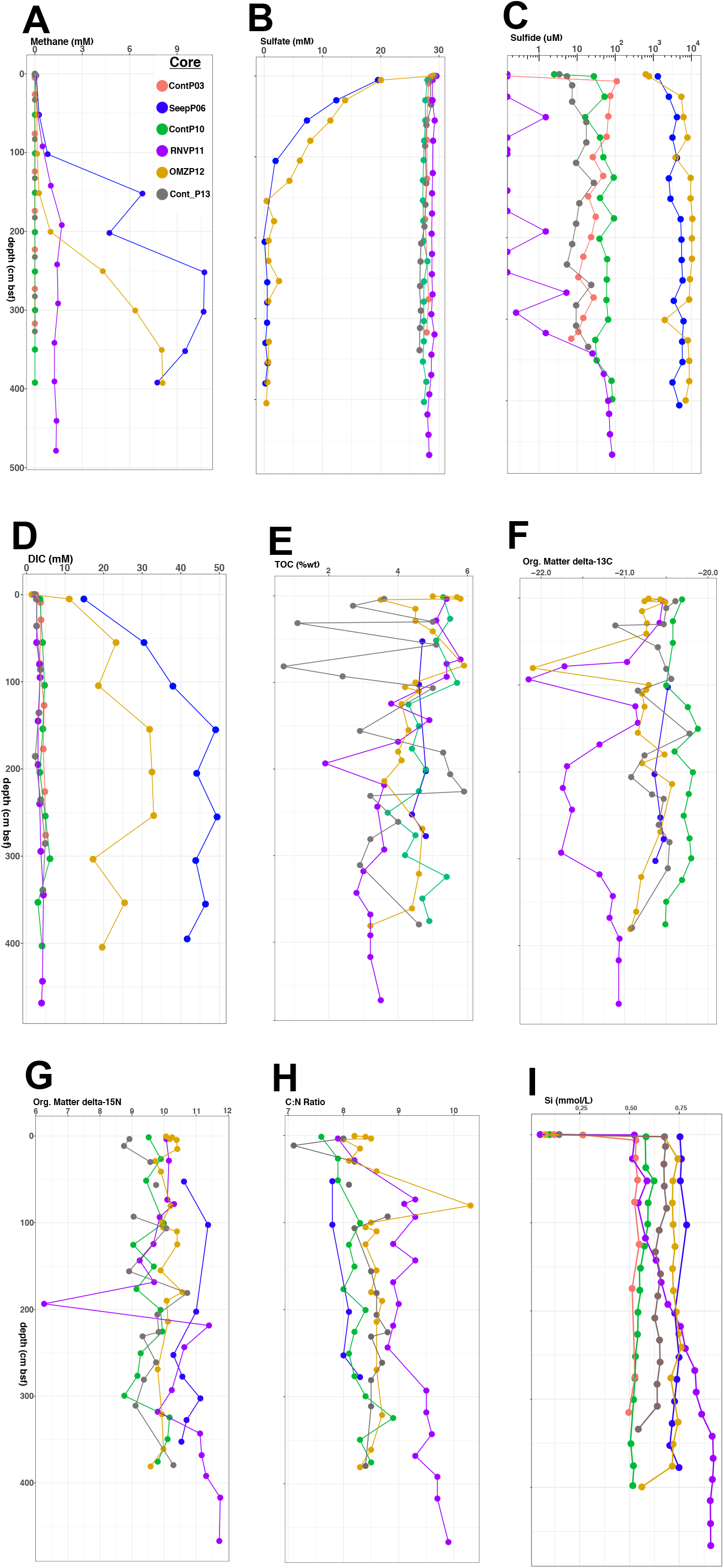
Geochemical profiles for A) Methane, B) Sulfate, C) Sulfide, and D) DIC porewater concentrations; E) Organic Carbon content in weight %; F) Organic Matter δ^13^C values, and G) δ^15^N values; H) Carbon to Nitrogen ratios, and I) Silica porewater concentrations. Geochemical data for site ContP03 are not available for the analyses depicted in panels E-H.

Core SeepP06 contains sulfide in millimolar concentrations throughout the core. Below the zone of sulfate-dependent methane oxidation at 1 m depth, methane accumulated to the highest concentrations of this survey, > 10 mM. Porewater DIC concentrations were consistently high and increased from 15 mM near the interface to nearly 50 mM with depth. These methane and DIC concentrations reached and in part exceeded the highest concentrations previously measured in Sonora Margin seep fluids (Paull et al., 2007). Sediments of core SeepP06 yielded only approximately half of the porewater volumes of other cores, indicating porewater loss by pressure-induced compaction. Thus, core SeepP06 represents sulfidic, methane- and DIC-soaked, compacted seep sediments from the pressure ridge aligned with the transform fault at the lower Sonora Margin (Simoneit et al., 1990;Paull et al., 2007).

Cores Cont03 and ContP10 share similar methane, sulfate, sulfide, and DIC profiles indicative of non-reducing conditions where sulfate-reducing and methanogenic activities remain minimal and biogenic sulfide and methane occur only in micromolar trace concentrations. With TOC between 4 and 6 wt %, the sediments of core ContP10 are organic rich and represent the hemipelagic seafloor sediments of Guaymas Basin that receive ample biogenic sedimentation, mostly by diatoms (Calvert, 1966). The δ^13^C values ranging from −20.12 to −20.51‰ are consistent with sedimentary organic material resulting predominantly from phytoplankton input (Teske et al., 2002). Slowly increasing δ^15^N values ranging from 9.04 to 10.17‰, and gradually increasing C:N ratios downcore are consistent with microbial utilization of nitrogen compounds in sedimentary biomass.

Contrasting with nearby core Cont10, Core RNVP11 shows the biogeochemical signatures of seawater inmixing and previous hydrothermal alteration at Ringvent (Teske et al., 2019). Subsurface-derived porewater methane in high concentrations of 1 to 1.5 mM coexists with porewater sulfate near seawater levels; sulfide is largely absent and reaches 10 to 100 μM only below 3 m depth. Core RNVP11 also stands out by having the lowest DIC concentrations of all cores, approaching seawater DIC in the upper layers. Below 1 mbsf, organic carbon δ^13^C values are the lowest for all cores, whereas C/N ratios are the highest, suggesting the influence of isotopically light and nitrogen-depleted fossil carbon sources (Figure 2). In contrast to other cores, Core RNVP11 shows a strong gradient of dissolved silica, increasing from 0.5 to 0.6 mM at the surface (similar to ContP10) towards 0.9 to 1 mM at the bottom (Figure 2 I). Silica dissolution is considered a marker of hydrothermal activity (Peter and Scott, 1988), and leads to elevated concentrations of dissolved silica in the water column of Guaymas Basin (Campbell and Gieskes, 1984).

Core OMZP12 differs from all other cores by its location in the oxygen minimum zone on the Sonora Margin slope (Calvert, 1964). Bottom water anoxia allows millimolar concentrations of porewater sulfide to permeate the entire sediment core including the surface, otherwise only seen in core SeepP06. The sulfate-methane transition zone occurs at approximately 1 and 2m depth for SeepP06 and OMZP12, respectively. Similar to core SeepP06, porewater DIC increases rapidly with depth, with a maximum value of 33mM at 254 cmbsf. Sediment TOC, δ^13^C, δ^15^N, and C:N ratio values generally resemble those of other cores in this survey.

Core ContP13, collected on the southeastern flanking region, differs from other cores by terrestrial input from the Yaqui River. Methane, sulfate, sulfide and DIC concentrations for this core follow similar depth profiles as observed for cores ContP03 and ContP10. However, TOC varies between ~1 and ~5 wt % in the first meter of sediment and between ~3 and ~6 wt % below, suggesting sedimentation pulses of varying organic carbon load. Sediment organic matter δ^13^C, δ^15^N and C:N ratios fall within the range of values observed for other cores in this survey.

### 2. Diversity of the Guaymas Basin archaeome

Rarefaction curves are plotted separately for samples in approx. 1 meter depth intervals to examine potential downcore trends (Figure 3). Starting at 3 m depth, observed species richness based on rarefaction summaries are lower in Ringvent (Core RNVP11) sediment compared to other sediments (Figure 3, Table S2). Substantially more sequence reads, and thus a higher number of observed species, were recovered from the Sonora Margin OMZ sediment (Core OMZP12) relative to the other surveyed sites, below 1 m depth (Figure 3 C-E). To account for different sequencing depths without resorting to rarefying the dataset (McMurdie and Holmes, 2014), we estimated total diversity using a non-linear regression model for ratios of consecutive frequency counts, a state-of-the-art method addressing the issue of heterologous sequencing depths affecting richness estimates (Willis and Bunge, 2015). Results from this model indicate no statistically significant (P_val_ < 0.05) differences amongst surveyed sites (Figure S1).

**Figure 3:**
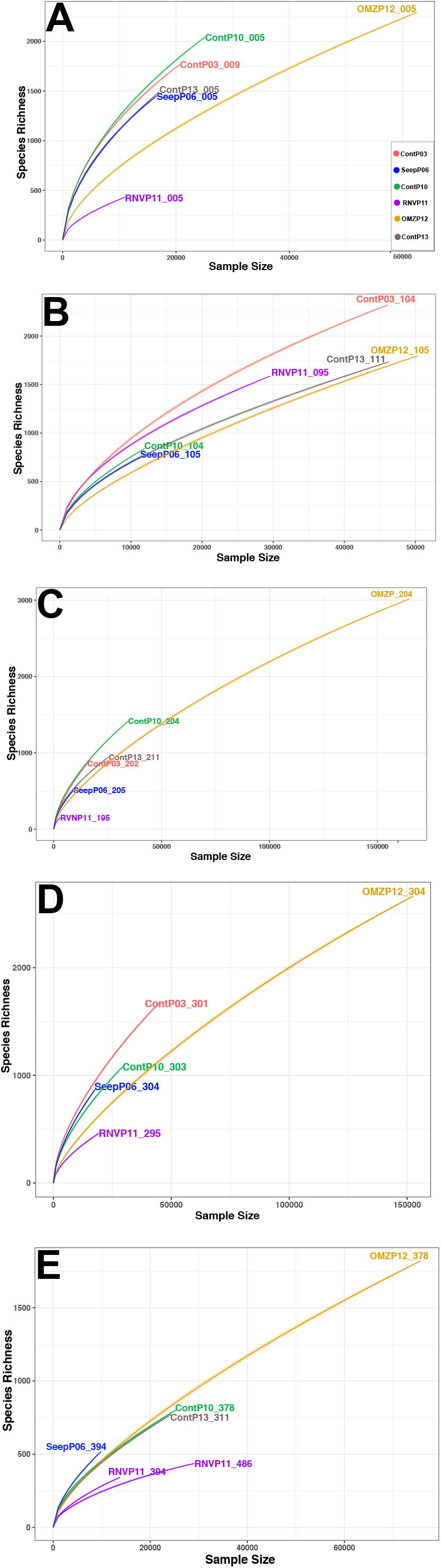
Depth mapped rarefaction summaries (color coded to match surveyed sites) for complete high quality sequence dataset depicting richness as number of OTUs (97% similarity clustered) observed per sequences sampled. A) samples near the interface (0-10cmbsf), B) samples from ~100cmbsf, C) samples from ~200cmbsf, D) samples from ~300cmbsf, E) samples from depths greater than or approximately equal 400cmbsf.

When beta diversity of the archaeal communities was examined for correlations with environmental metadata using two-dimensional principal coordinate analysis, distinct clustering patterns are observed (Figure 4A). Surface communities for all cores except OMZP12 tightly cluster along negative axis 1 values. All OMZP12 samples cluster along axis 1 values greater than 0.01 independently of sediment depth. Separation along axis 2 partitions the SeepP6, RNVP11, and OMZP12 cores (with positive axis 2 values), from subsurface samples of control cores ContP3, ContP10, and ContP13 with negative axis 2 values; the surface samples of these cores cluster separately (Figure 4A). The influence of environmental factors (*i.e.*: methane, sulfate, sulfide, DIC, water depth, and sediment depth) on community ordination is complex (Figure 4B-G), and it appears likely that clustering patterns are not driven by these environmental parameters alone. Water column depth (Figure 4F) appears to drive core OMZP12 clustering along larger positive values for axis 1, but most likely represents a proxy for the influence of the oxygen minimum zone at this depth.

**Figure 4:**
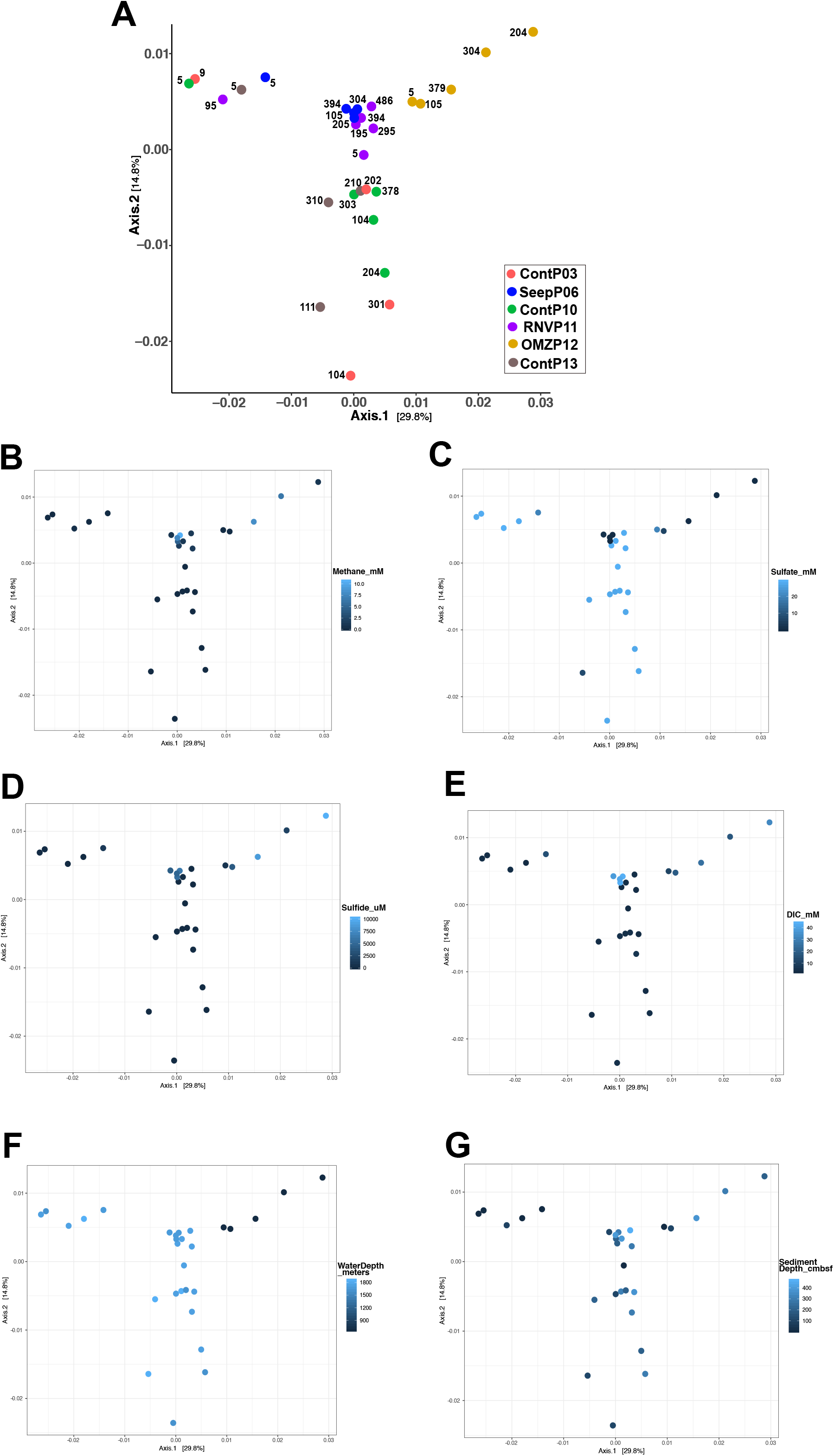
A) Two-dimensional Principal Coordinate Analyses of Bray-Curtis dissimilarity distances from r-log normalized sequence count data. Each community plotted is color-coded to the core site and numerical labels indicate sediment depth (cmbsf). The first and second axes explain 29.8 and 14.8% of the variance, respectively. Environmental metadata superimposed on ordination plot are B) methane, C) sulfate, D) sulfide, and E) DIC concentrations, F) water depth, and G) sediment depth.

### 3. Network analysis

Network analysis based on the co-occurrence of all OTUs in each sample reveals that the deepest communities recovered from Core RNVP11 exhibit the greatest degree of separation (Figure 5). At a maximum ecological (Bray-Curtis) distance of 0.8 (*i.e.*: the maximum distance allowed between two samples to be considered connected in the graphical model), most samples share taxa co-occurrence patterns, except for the deepest communities from core RNVP11 (Figure 5A). Decreasing the minimum ecological distance in the model to 0.5, resolves three independent network clusters (Figure 5B). Here, the two deepest samples from core RNVP11 share similar taxa co-occurrence patterns only with each other and are excluded from the two emergent additional networks. In one of these networks, communities near the seawater interface of all cores, with the exception of core RNVP11, connect at no more than 1-degree of separation. Interface sample SeepP06 5cmbsf connects the near-interface sample cluster to all core SeepP6 subseafloor (depth > 1mbsf) samples. A third independent network shows non-random taxa co-occurrence amongst subseafloor control sediments (cores ContP3, ContP10, ContP13), a subseafloor and a near-interface sample from core RNVP11, and all samples from core OMZP12; the three deepest OMZP12 samples are only peripherally connected (Figure 5B).

**Figure 5.**
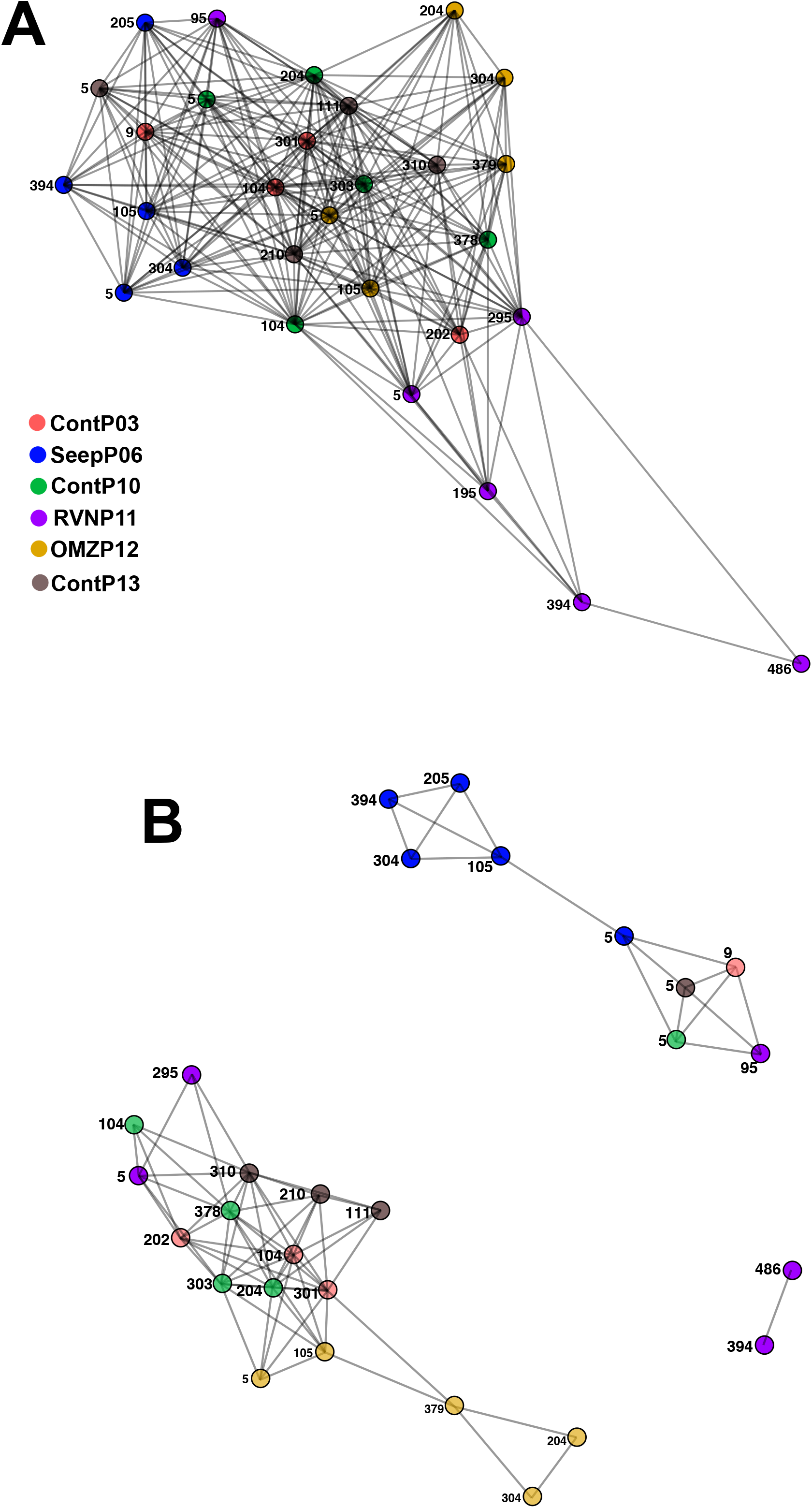
Network analysis based on the co-occurrence of all OTUs at in each sample. Nodes represent all archaeal communities analyzed in this study. Nodes are color-coded to match descriptions from Figure 1A. Edges are unweighted interactions depicting OTU co-occurrence meeting arbitrary thresholds. A) Co-occurrence network threshold set at a maximum Bray-Curtis distance of 0.8. B) Co-occurrence network threshold set at a maximum Bray-Curtis distance of 0.5.

### 4. Community composition

Class-level community descriptions (SILVA132) assigned large membership fractions of the archaeal communities to the Bathyarchaeia, Hadesarchaeaeota, and Thermoplasmata (Figure 6). The Class Methanomicrobia, comprising methane-producing and methane-oxidizing members of the Euryarchaeota, was detected in multiple cores, but appeared most frequently at depth in core SeepP06. An in-depth summary of the Methanomicrobia reveals the presence of methanogenic families (eg: Methanomicrobiaceae) and anaerobic, methane-oxidizing ANME lineages (ANME-1, various ANME-2). Notably, ANME-1 archaea dominate core SeepP06 sequence assignments comprising nearly 40% of the total community at 394 cmbsf in this core (Figure S2, Table S3). Order- or higher-level community taxonomic descriptions for all samples generally contained 60% or greater unclassified community fractions (data not shown) when automated taxonomic assignments were performed. In order to not rely on the uncertain output of taxonomy pipelines, and to resolve archaeal taxonomy assignments in a manner that is consistent with broadly accepted usage (Spang et al., 2017), we also describe community composition based on phylogenetic placement of dominant sequence variants for the most numerous 25 Operational Taxonomic Units (OTUs).

**Figure 6:**
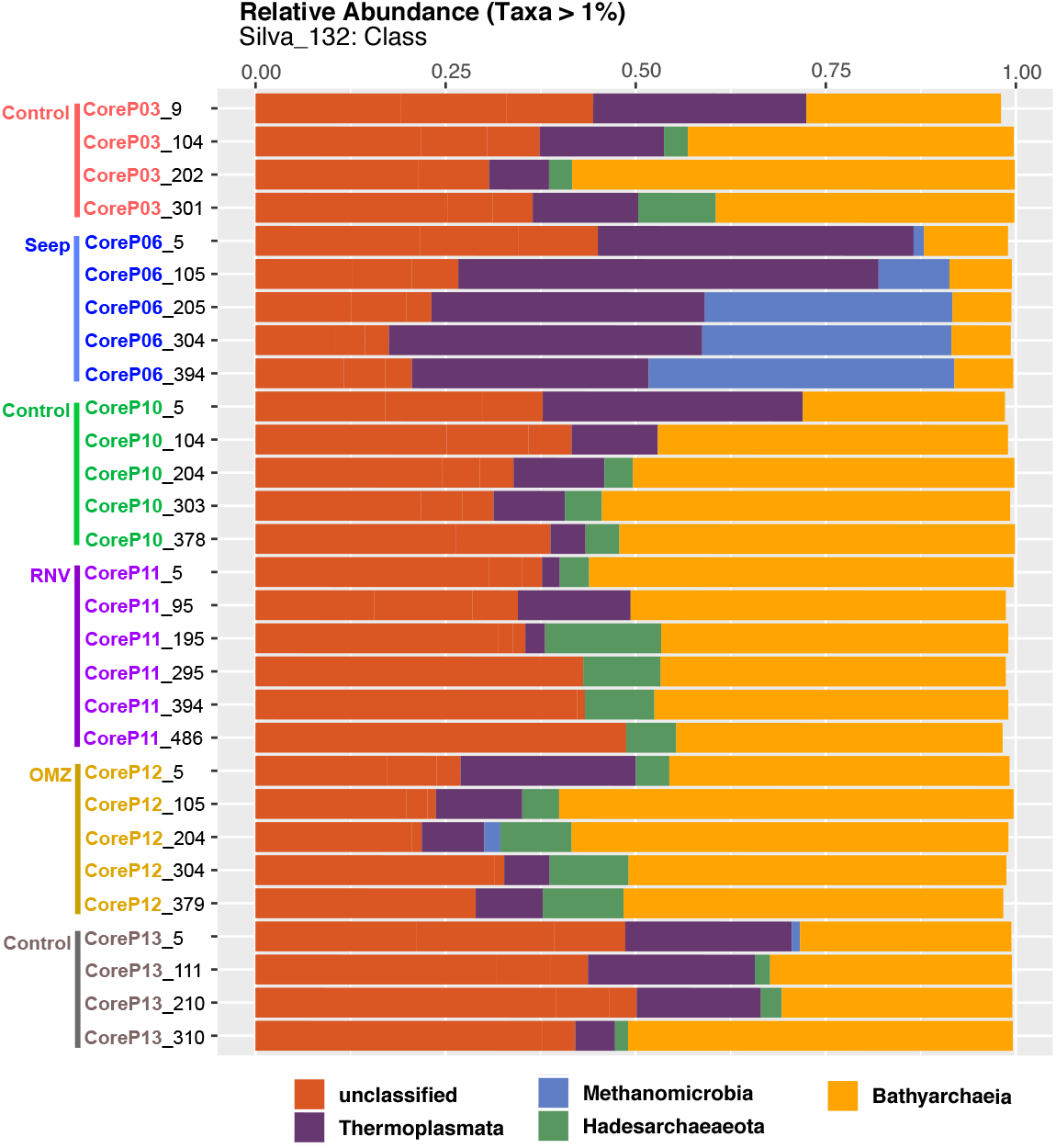
Class-level community composition of all depths for all cores in this study. Core labels are color-coded to match the collection sites depicted in the bathymetric map in Figure 1.

The majority of high-quality sequences in this study (73.0%) clustered into 25 OTU lineages (Figure 7A). Archaeal communities were largely dominated by OTU lineages related to the Bathyarchaea (16 out of the top 25 OTUs). OTUs 01 to 03, the three most abundant lineages, belong to the MCG-1, MCG-2, and MCG-3 Bathyarchaea subgroups (Kubo et al., 2012) respectively, with close relatives recovered from Guaymas Basin and globally-dispersed subseafloor habitats (Figure 7B). High abundance lineages related to the Marine Benthic Group-D within the Thermoplasmata (MBG-D; OTUs 04, 10, 11, 21, and 23) and the Terrestrial Miscellaneous Euryarchaea Group (TMEG; OTU 08) were recovered from all cores and core depths except the subsurface of Ringvent (Core RNVP11, depth > 1mbsf). Two highly abundant lineages represented by OTUs 05 and 18 (Figure 7C) were recovered from every core except core SeepP06, and identified as relatives of the Hadesarchaea, formerly known as South African Gold Mine Euryarchaea Group (SAGMEG)-1 (Baker et al., 2016). Lastly, OTU 14 clustered within anaerobic ANME-1 methanotrophs and was most closely related to ANME-1 phylotypes from cold seep, hydrate and brine habitats; OTU 14 did not affiliate with the thermophilic ANME-1Guaymas lineage recovered from hydrothermally active, hot sediments in Guaymas Basin (Biddle et al., 2012) (Figure 7C).

**Figure 7:**
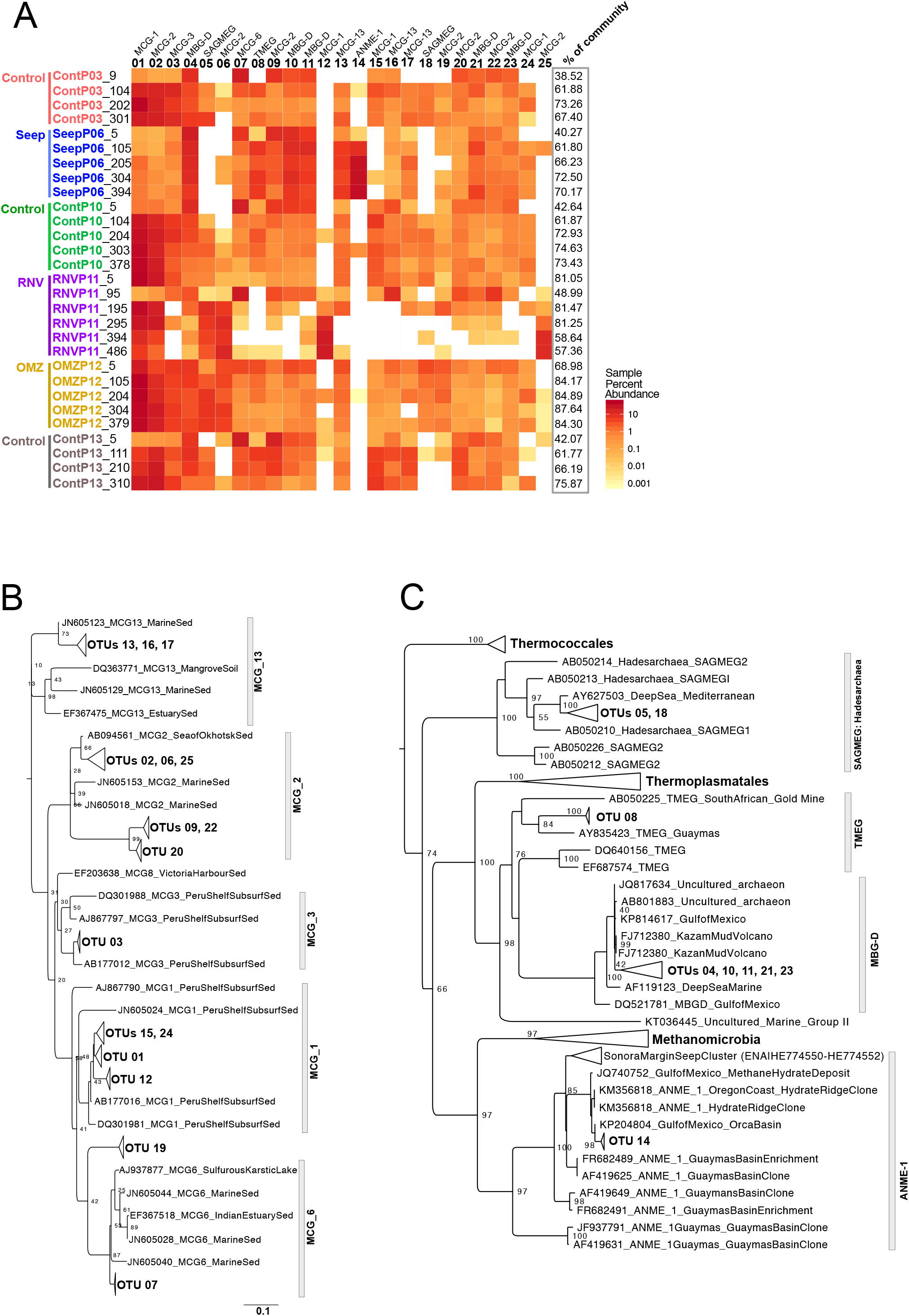
A) Heatmap depicting percent abundance distribution for the 25 most abundant OTUs, representing 73.0% of all high-quality sequences in this study, for all cores and across all depths. Core labels are color-coded to match the collection sites depicted in the bathymetric map in Figure 1A. The phylogenetic association of each OTU lineage is depicted above each OTU header. The percent of total reads represented by the 25 most abundant OTUs in each community is shown in the column labeled “% of community”. Maximum likelihood phylogenetic trees, with 100 bootstrap support, placing the top 25 most abundant OTUs within the following lineages: B) Bathyarchaea, C) the Euryarchaeotal lineages MBG-D, TMEG, SAGMEG, and ANMEs.

### 5. Differential taxon abundance estimations across ecological niches

Differential abundance analyses (Wald Test, Pval = 0.01) were performed on various ecological-models following potential environmental niches suggested by ordination patterns (Figure 8). Only OTU lineages amongst comparison groups containing more than 100 sequences were used for each test. We tested the influence of sediment depth in the absence of seepage or hydrothermal influence, the impact of the oxygen minimum zone waters on surficial and subsurface sediments, and the effect of hydrothermal disturbance. These analyses have to be qualified by the fact that they are based on patterns of sequence frequencies, which are derived from the archaeal community but do not necessarily represent it in identical proportions.

**Figure 8:**
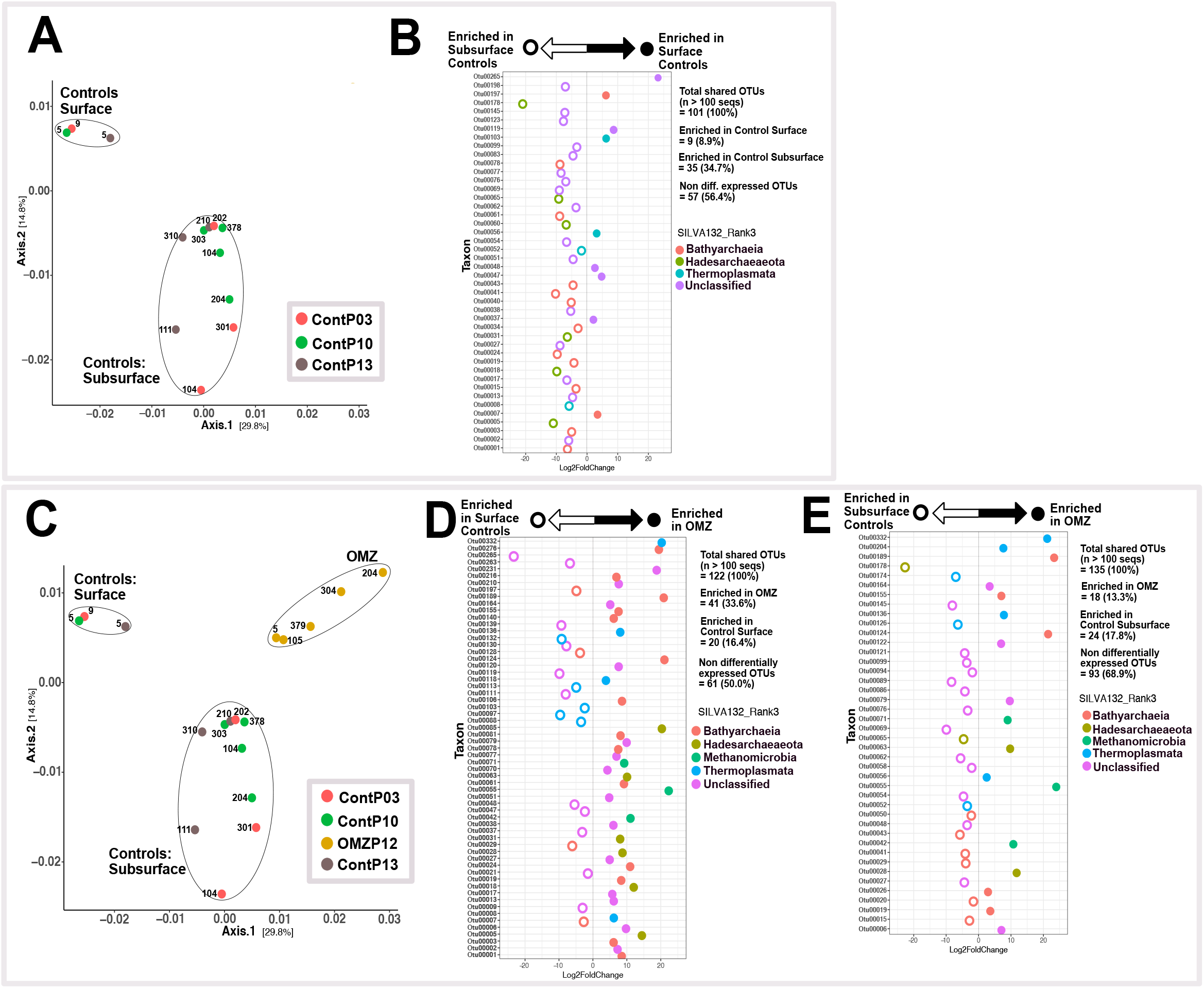
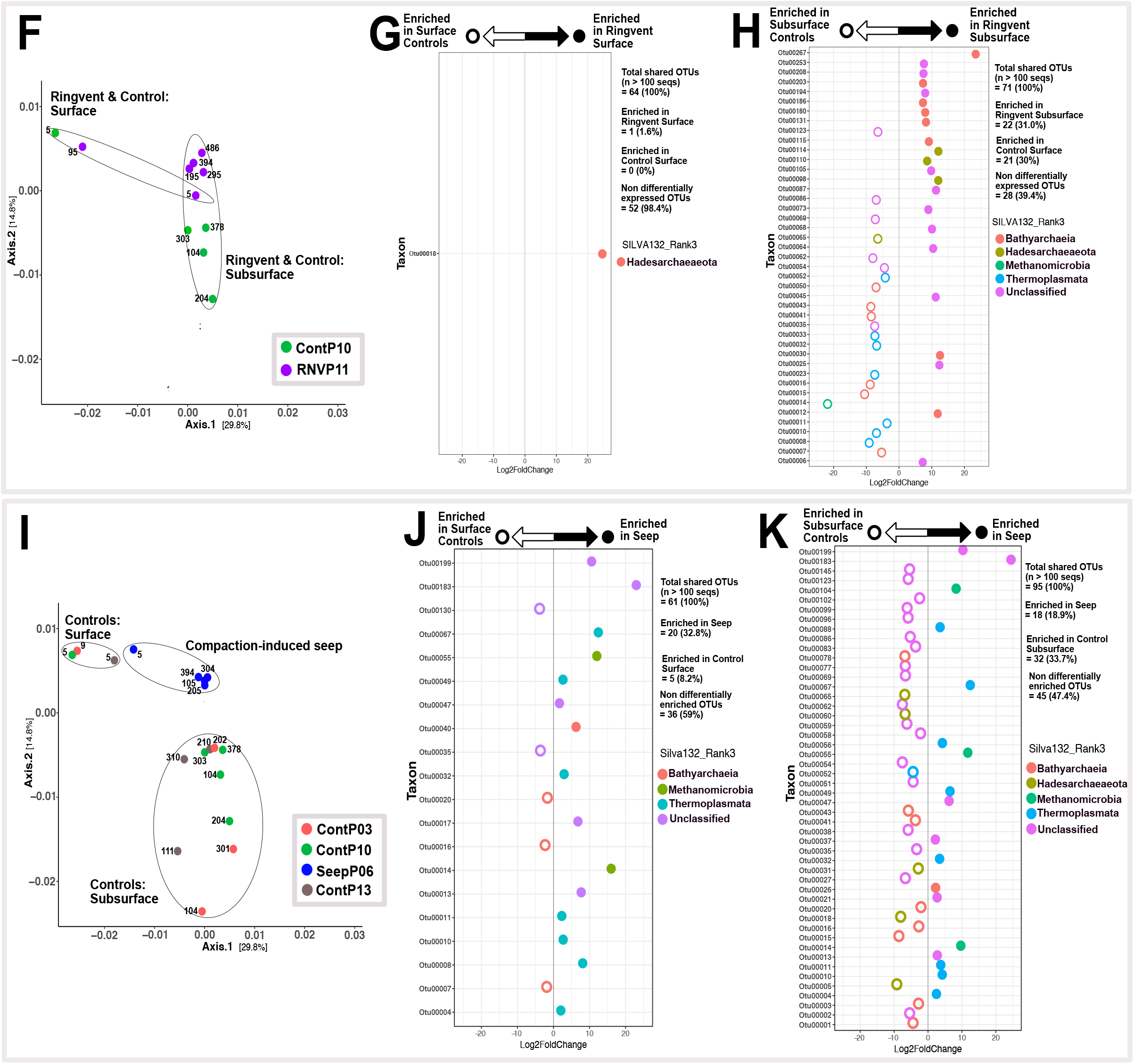
Differential Abundance Analyses bases on Wald’s test (significance: alpha = 0.01). A) Ordination depicting archaeal community clustering for surface and subsurface samples of control sites ContP3, ContP10, and ContP13. B) Differentially abundant OTUs in near-surface versus subsurface communities from control sites. C) Ordination depicting community clustering in OMZP12, and surficial vs. subseafloor control sites. D) Differentially abundant OTUs in OMZP12 compared to surficial and E) subsurface communities from control sites. F) Ordination depicting community clustering in RNVP11 to Cont10. Differentially abundant OTUs in RNVP11 and Cont10 for G) surface samples and H) subsurface samples. I) Ordination depicting community clustering in SeepP06, and surficial and subseafloor control sites. Differentially abundant OTUs in SeepP06 samples compared to J) surficial and K) subsurface communities from control sites. Note: OTUs 13, 17 and 21 are color-coded as “unclassified” by SILVA132, but the phylogeny identifies them as members of MCG-13 (OTUs 13 and 17) and MBG-D (OTU 21).

To test the estimated differential abundance of archaeal community members in near-surface (depth < 1mbsf) and subsurface (depth > 1mbsf) communities under conditions of normal hemipelagic sedimentation, we selected cores ContP3, ContP10, and ContP13; these cores lack hydrothermal, seepage or OMZ influence and, therefore, show the archaeal community of organic-rich Guaymas Basin sediments in the absence of these selective factors. Here, over 43% of OTUs (n > 100 seqs) are differentially enriched with depth (Figure 8A & B). Most differentially abundant taxa are estimated to be significantly less abundant in near-surface relative to subsurface control sediment (Figure 8B). Subsurface enriched lineages include members of various Bathyarchaeal groups (MCG-1, MCG-3, and MCG-6), a TMEG lineage (OTU 08) closely related to clones previously recovered from Guaymas Basin, and a Hadesarchaea lineage (OTU 18) whose closest relatives are clones from deep Mediterranean waters (Figures 7B & C). Among the top 25 OTUs in this study, the only highly abundant lineage that is significantly (Pval = 0.01) enriched in near-surface relative to subsurface control sediment (Figure 8A), is OTU 07, related to Guaymas Basin and Indian estuary sediment MCG-6 clones (Figure 7B). The same lineage was also found to occur preferentially in surficial estuarine sediments in the White Oak River, while avoiding the sulfide-rich, sulfate-reducing and methanogenic conditions just a few centimeters downcore (Lazar et al., 2015).

When archaeal abundance in all oxygen minimum core OMZP12 samples was checked against the shallow sediment samples of the control cores (ContP03, ContP10, ContP13, see Figure 8C and D), over 50% of archaeal OTUs (n > 100 seqs) present in surficial controls were differentially enriched in core OMZP12. Of the 25 most highly abundant OTUs in this study, those that were differentially abundant in this comparison (14 OTUs) were predominantly enriched in core OMZP12 (eleven of fourteen high abundance OTUs, Figure 8D). Only OTUs 07, 09, and 21, representing the Bathyarchaeal groups MGC-6 and MCG-2, and a MBG-D lineage, respectively, were enriched in surficial control sediment (Figure 8D).

We also tested for differentially abundant taxa between all core OMZP12 samples and subsurface control sediment (Figure 8E). Here, only 31.1% of OTUs (n > 100 seqs) were differentially enriched; of the 25 most abundant OTUs in this study only 4 were differentially enriched. Of these four differentially enriched archaeal OTUs, OTUs 15 and 20, Bathyarchaeal lineages in the MCG-1 and MCG-2, respectively, were enriched in subsurface control sediment. OTU 06 within the MCG-2 group, and OTU 19, an MCG lineage tenuously related (bootstrap value < 70%) to MCG-6, were enriched in core OMZP12. Overall, archaeal types occurring in anoxic subsurface sediments of core OMZP12 resemble those in other subsurface cores to a large extent.

The hydrothermally influenced Ringvent core RNVP11 was compared against control core ContP10, located only 1.6 km further west, at the same depth and local sedimentation regime (Figure 8F-H). In surficial sediment (> 1 mbsf) this comparison revealed only one differentially enriched phylotype, OTU 18 within the Hadesarchaea (Figure 8G). On the other hand, the comparison of subsurface (>1 mbsf) communities identified 43 OTUs, or 61% of all shared OTUs (n > 100 seqs), as differentially abundant between these two sites (Figure 8H). Focusing on the top 25 highest abundance OTUs in this study, eleven OTUs were differentially enriched. Eight taxa, comprised of Bathyarchaeal, TMEG, MBG-D, and ANME-1 representatives, were significantly enriched in subsurface control sediment relative to Ringvent subsurface sediment (OTUs: 07, 08, 10, 11, 14, 15, 16, and 23). The remaining three taxa (OTUs 06, 12, and 25), enriched in the Ringvent subsurface, were classified as Bathyarchaea related to the MCG-1 and MCG-2 subgroups.

Core SeepP06, from compacted seep sediments, was checked against both shallow and subsurface sediment samples of the control cores (ContP03, ContP10, ContP13, Figure 7I-K). 41% of archaeal OTUs (n > 100 seqs) present in shallow controls and seep sediment were differentially enriched (Figure 8J). Of the 25 most highly abundant OTUs, most were predominantly enriched in seep sediment and included lineages classified as ANME-1, MBG-D, and TMEG (Figure 8J). When comparing differentially abundant taxa between core SeepP06 and subsurface control sediment, almost 53% of archaeal OTUs (n > 100 seqs) were differentially enriched (Figure 8K). Most of the OTUs in this comparison, including the most abundant OTUs in this study (OTUs 01-03), were enriched in control subseafloor sediments rather than in core SeepP06 sediment (Figure 8K).

## Discussion

### 1. Complex determinants of archaeal ecosystem structure

Overall, complex physical and geochemical factors structure sedimentary habitats and depth-related niches for archaea in Guaymas Basin. Archaeal community ordination patterns reveal niche differentiation and some unexpected clustering patterns among the different sites (Figure 4A).

Most notably, surficial communities of background control sediments ContP03, ContP10, and ContP13, at 5 to 10 cm depth, cluster away from their respective subsurface communities near 1 m depth and below (Figure 4A), implying an ecological bifurcation into two distinct niches. Taxa co-occurrence network patterns support this differentiation between shallow and subseafloor control sediment sites (Figure 5B). Possibly, the availability of electron acceptors such as oxygen, nitrate or oxidized metals drives the depth-dependent niche separation observed in background control sediment sites (ContP03, ContP10, and ContP13). Surface archaeal communities continually change as sediment layers accumulate; given high sedimentation rates of 0.23 to 1 mm/y at Guaymas Basin (Teske et al., 2019), it takes approx. 1000 to 4000 years for background surface communities to transition to subsurface communities at 1 m depth. Interestingly, shallow vs subsurface differentiation is less apparent in seep, OMZ or hydrothermally influenced sites (Figure 5B); parameters other than sediment depth or surficial redox regime are shaping archaeal community composition in seepage- or hydrothermally-influenced habitats, compared to the control sites.

The cores SeepP06 and RNVP11 represent different geochemical regimes (compaction-induced continental margin seepage vs hydrothermal circulation, respectively), yet these two sites cluster tightly in ordination space (Figure 4). Individual geochemical factors, for example, the sulfidic, methane- and DIC-rich conditions in SeepP06 would have indicated that OMZP12 should be its closest equivalent (Figure 2). The unexpected clustering of SeepP06 and RNVP11 suggests that factors beyond current geochemical conditions, for example recent environmental disturbance, can influence archaeal community structure. At Ringvent (RNVP11), sedimentary community diversity may have been reduced during prior episodes of thermal purging or high methane flux (Teske et al., 2019), selecting for a resilient, yet potentially less diverse (Figure 3), “survivor” community.

Lastly, community ordination differentiates OMZ sediment from all other sedimentary habitats (Figure 4). While water depth appears to have a strong influence on OMZ sediment ordination (Figure 4F), we propose that differences in redox potential at the sediment interface due to its direct contact with oxygen-depleted water (Calvert, 1964) rather than water column depth, is the key environmental constraint driving the ordination patterns of OMZ sediment.

### 2. A “forest view” of Archaea in Guaymas Basin sediments

Archaea observed in this sedimentary habitat survey belong to the Bathyarchaea, the MBG-D and TMEG lineages within the Thermoplasmatales, the Hadesarchaea (SAGMEG), and ANME-1 lineages, as shown previously in a sequencing survey using different archaeal primers (Teske et al. 2019). The uncultured Bathyarchaea and MBG-D archaea pronouncedly dominate the dataset, and cold anoxic marine sediments globally (Kubo et al., 2012;Lloyd et al., 2013). Bathyarchaea play an important role, tantamount to that of the domain Bacteria, in the remineralization of complex organic matter in marine sediment (Lloyd et al., 2013); some of their members (MCG-8 lineage) use lignin, the second most common biopolymer on Earth, as an energy source (Yu et al., 2018). Since Bathyarchaeota harbor the Woods-Ljungdahl pathway, they are implied in acetogenic subsurface metabolism (He et al., 2016), but it remains open whether acetogenic pathways are used for net autotrophy, or derive their substrates from organic carbon sources (Lever et al., 2010); similar considerations apply to the metabolically versatile MBG-D archaea (Zhou et al., 2019). Some Bathyarchaea harbor genes of the MCR complex, suggesting methylotrophic methanogenic activity and, perhaps, syntrophic interactions with sulfate reducing bacteria leading to the anaerobic oxidation of methane (Evans et al., 2015). However, the dominant Bathyarchaea OTUs recovered in this study (MCGs 1, 2, 3, 6, and 13, Figure 7B) are only distant relatives of methane-cycling marine Bathyarchaea, which fall into MCGs 15 and 16 (Evans et al., 2015). Hadesarchaea, originally described as the South-African Gold Mine Miscellaneous Euryarchaeal Group (SAGMEG), are metabolically versatile anaerobic heterotrophs with the metabolic potential for CO and H2 oxidation coupled with nitrite reduction to ammonia and are found in environments across broad (4 to 80°C) thermal ranges (Baker et al., 2016). One of two frequently recovered Hadesarchaea lineages (OTU 05) is conspicuously enriched in subsurface sediment in Ringvent (RNVP11), where low observed sequence richness (Figure 3) coincides with evidence (high silica porewater concentrations at depth indicative of a thermal dissolution of sedimentary diatoms, Figure 2I) for a thermal purge in the past (Teske et al., 2019).

Unsurprisingly, the methane-cycling Methanomicrobia are rare in background sediments (ContP03, Cont10, and Cont13) but are strongly enriched in core SeepP6 and, to a much lower extent (slightly over 2%), in core OMZP12 at 204 cm depth (Figures 6 and S2). Interestingly, a single ANME-1 OTU lineage, OTU14, is highly enriched in SeepP06 sediments (Figure 7A and C). This lineage is closely related to ANMEs recovered from cold, anoxic habitats, such as seafloor seep sediments, methane hydrates, and hypersaline anoxic basins, and distinct from previously described ANME-1 phylotypes from Sonora Margin cold seep sediments and potentially thermophilic ANME-1 phylotypes from Guaymas Basin hydrothermal sediments (Holler et al., 2011;Biddle et al., 2012). Although ANME-1 archaea were generally assumed to be obligate methanotrophs, this assumption has been challenged and this lineage has been proposed as potentially methanogenic, based on its occurrence and activity in sulfate-depleted sediments (Lloyd et al., 2011;Kevorkian et al., 2020); thus, the biogeochemical role of these archaea would be modulated by the presence or absence of sulfate, or concomitant changes in electron donors. ANME-2 and cultured methanogenic lineages were observed in low percent abundances in all cores in this study (Figure S2). Interestingly, ANME-2 lineages were extremely rare, representing less than 0.05% of any sample and less than 0.02% of any SeepP6 community (Figure S2, Table S3). The prevalence of ANME-1 over ANME-2 in the El Puma cores is consistent with the ecophysiological preference of ANME-1 archaea for reducing, sulfidic subsurface sediments, and the preference of ANME-2 for near-surface sediments with intermittently oxidizing conditions (Rossel et al., 2011;Ruff et al., 2015). Previous surveys of mat-covered seep sediments on the Sonora Margin have revealed transitions from ANME-2 towards ANME-1 within short push cores of max. 17 cm depth (Vigneron et al., 2013).

Overall, we hypothesize that the Archaeome in the sedimented flanking regions of Guaymas Basin is generally fueled by heterotrophic processes including the degradation of proteins, polymeric carbohydrates (Ziervogel and Arnosti, 2020), and accumulating lipids (Teske et al., 2002) resulting from high sedimentation rates driven by high levels of primary production in the water column. Diverse niche communities allow the Guaymas Archaeome to adapt to environmental challenges, such as hydrothermalism or methane seepage, that are common in the greater Guaymas Basin area; thus, the site-specific complexity of the Guaymas Archaeome underpins its resilience.

### 3. Ecological comparisons: Differentially abundant taxa across sedimentary habitats

#### 3.1 Near-surface vs. subsurface sediment niches

When comparing the surficial to the subsurface archaeal populations in background control sediments, the majority of OTUs estimated to be differentially abundant are significantly more enriched in the subsurface relative to the surficial sediment, suggesting that benthic archaea prefer subsurface conditions (Figure 8A, B). This trend may also reflect the impact of electron acceptors; for example, oxygen permeates background sediments in Guaymas Basin for at least one centimeter [(Teske et al., 2016), Figure 8B therein]. Following a recently proposed model for benthic microbial communities (Starnawski et al., 2017) the archaeal community at the oxic water-sediment interface likely undergoes downcore selection, based on site-specific selective pressure, resulting in reduced diversity with depth but a higher prevalence of subsurface-adapted taxa within a few thousand years after burial in anoxic subseafloor sediment. Benthic archaea, predominantly Bathyarchaea and MBG-D lineages, survive on residual carbon sources that remain after burial and microbial degradation in surficial sediments (Lloyd et al., 2013). Interestingly, catabolic activity and electron donor diversity, rather than terminal electron acceptor type or burial time, appear to drive bacterial OTU richness in anoxic subseafloor sediment (Walsh et al., 2016). This niche construction mechanism, driven by the biotic microenvironment as opposed to abiotic environmental filtering (Aguilar-Trigueros et al., 2017), is potentially widespread across the large habitable volume represented by non-hydrothermally influenced subseafloor sediments in Guaymas Basin.

#### 3.2 OMZ vs. control sediment

When comparing OMZ and surficial background control sedimentary communities, 50% of high abundance OTUs found across both habitats are estimated to be differentially enriched (Figure 8D). Two thirds of the differentially enriched taxa were enriched in the OMZ rather than the surficial background sediments. The MCG lineages MGC-6 and MCG-2 (OTUs 9 and 7) and a MBG-D phylotype (OTU 21) are enriched in the surficial background controls relative to the OMZ sediment (Figure 8D). Interestingly, MCG-6 members bear hydrolases that specifically target plant-derived polymeric carbohydrates (Lazar et al., 2016), a potential trait-environment relationship that may differentiate surficial background control from OMZ sediment communities (Figure 8D). When comparing subsurface background control and OMZ sediment communities, the number of OTUs estimated to be differentially enriched was about equal across both environments; however, the majority (68.9%) of high abundance OTUs show no significant differences in their estimated abundances at all (Figure 8E). This implies that the subsurface, rather than surficial, background control communities are more similar to the OMZ communities, a point also corroborated by taxa co-occurrence network analysis (Figure 5B). Thus, oxygen depletion in background subsurface sediment, and oxygen depletion through the overlying oxygen minimum zone of the water column (Calvert, 1964), result in some convergence between archaeal communities across geographically distant and environmentally distinct sedimentary habitats.

#### 3.3 Ringvent vs. control sediment

The surficial archaeal communities of Ringvent (RNVP11) and its nearby control site (ContP10) are similar to each other, as indicated by extensive co-occurrence networks (Figure 5) and by the lack of differential enrichment between the two cores (Figure 8G). A member of the Hadesarchaea, OTU18, is estimated to be significantly enriched in Ringvent surficial sediment relative to the control; otherwise differences in taxon abundance across these habitats are negligible. Both sites are only 1.6 km apart and therefore most likely share recent depositional histories and microbial inoculum sources, which validates core ContP10 as a site-specific control for assessing the environmental determinants structuring subsurface archaeal communities at Ringvent. The reduction in sequence recovery and, potentially, archaeal community richness in subsurface Ringvent (RNVP11) sediment (Figure 3) is attributed to environmental selection via hydrothermal purging or methane seepage driven by recent sill emplacement that continues to drive hydrothermal circulation, selecting against microbes unable to withstand chemical or thermal changes associated with hydrothermal circulation (Teske et al., 2019). Thus, OTUs may be enriched in Ringvent subsurface sediment relative to its nearby control site (Figure 8H) via two possible ecological scenarios; i) surviving resilient microbes could dominate the habitat after their competitors have been removed, and ii) new arrivals after the disturbance could efficiently recolonize the depopulated surface sediment.

#### 3.4 Seep vs. control sediment

Differential abundance comparisons show that the ANME-1, MBG-D, and TMEG lineages are significantly enriched in the seep sediments, compared to controls (Figure 8I-K). Generally, methane seeps are specialized microbial benthic habitats where methanotrophic archaea (ANME) and syntrophic Deltaproteobacteria oxidize methane anaerobically exploiting sulfate as an electron acceptor (Lloyd et al., 2010;Ruff et al., 2015). The dominance of these inter-domain syntrophic partners distinguishes seafloor seep habitats (Ruff et al. 2015). Archaeal community structure in SeepP06 sediments differs little with depth; it is most similar, in terms of taxa overlap, to other samples from the same core (Figures 4 & 5). Therefore, the influence of cold seepage drives community selection to a greater degree than the environmental factors associated with depth-dependent niche differentiation observed in background control sediment.

Comparison with other Sonora Margin cores highlights the seep characteristics of core SeepP06. Based on the presence or absence of major archaeal lineages, SeepP06 archaeal communities are similar to surficial (<1mbsf) communities from Sonora Margin cold seeps, predominantly comprised of Thermoplasmata (MBG-D), Bathyarchaea, and ANME lineages (Cruaud et al., 2017). The SeepP06 archaeal communities share dominant archaeal lineages – the Thermoplasmatales (MBG-D), Lokiarchaeota and Bathyarchaeota – with Sonora Margin subsurface sediments [core BCK1, (Vigneron et al., 2014)]. Interestingly, the high proportion of ANME-1 archaea in SeepP06 is not shared by the Sonora Margin subsurface core (Vigneron et al., 2014). The Sonora Margin subsurface sediment core has a deeper methane/sulfate interface than Seep06, ca. 4-5 m instead of 1 m, and contains little sulfide above 5 m depth, indicating strongly attenuated seep influence in core BCK1 compared to SeepP06.

### 4. Core-specific features of the benthic Archaeome

Controls that structure microbial communities in hydrothermal sediments of Guaymas Basin have been studied extensively; for example, extreme temperature and porewater gradients shape microbial population structure, genomic repertoire and activities within a few centimeters depth beneath the seafloor (McKay et al., 2012;McKay et al., 2016;Dombrowski et al., 2018). However, ecological factors influencing microbial life in other sedimentary habitats at Guaymas Basin are comparatively unconstrained. By comparing archaeal communities in diverse sedimentary habitats to background controls representative of standard hemipelagic sedimentation, characteristic responses of the archaeal communities to these distinct environmental settings are becoming apparent. Compaction-induced seepage near the base of the Sonora Margin, and the resulting methane- and sulfide-rich porewater conditions in core SeepP06, selected for anaerobic methane-oxidizing archaea (ANME-1) and MBG-D archaea within the Thermoplasmata, and reduced the relative proportion of Hadesarchaea and Bathyarchaeota. Prior disturbances by hydrothermal impact or strong methane seepage, exemplified in the Ringvent sediments (RNVP11), also strongly differentiated sedimentary archaeal communities from those in background controls. Observed community richness in RNVP11 based on rarefaction curves are reduced throughout much of the core; these results resembled the outcome of a parallel study using different archaeal primers, and bacterial primers as well (Teske et al., 2019). Lastly, anoxic bottom waters impinging on the sediment on the upper Sonora Margin (OMZP12) drive similarities between anoxic surficial sediment at this site and anoxic subsurface background control sediments. The anoxic redox state of the water-sediment interface may also enhance archaeal richness estimates in the upper sediment column, potentially by facilitating the pelagic-benthic transition of archaea, or selecting against a bacteria-dominated interface (Xia et al., 2017). In brief, the archaeal communities of different cores respond in different ways to specific local controls.

### 5. Environmental history determines ecological context

The sediment cores shared similar biogeochemical parameters, such as sedimentary TOC, and organic matter δ^13^C, δ^15^N and C:N ratios. Repeatedly, studies of uncultured microbes in the sedimentary subsurface tried to correlate community composition with a wide range of biogeochemical or thermal parameters, in the hope that these linkages provide insights into habitat preference and ecophysiology of uncultured archaea (Durbin and Teske, 2012;Lazar et al., 2015;McKay et al., 2016). While this strategy can yield valuable results, we caution that patterns of archaeal community composition are not deterministically linked to biogeochemical parameters alone, rather, the full context of an ecological interpretation requires that biological and geochemical observations are integrated with the environmental setting and history of a site. For example, the lighter δ^13^C values of sedimentary organic matter in RNVP11 (trending towards −22 ‰ compared to most values clustering between 20 and 21 ‰), the slightly elevated C:N ratios at this site, increased Si concentrations at depth, or the elevated methane content superimposed on seawater-like porewater characteristics, are not in themselves critical factors that determine biological metrics in this core; these factors are significant because they reveal a depositional history of organic-rich sediments overprinted by relatively recent hydrothermalism and methane flux that has left its footprint on the present-day archaeal community. In another example, the archaeal communities of cores SeepP06 and OMZP12 would be assumed to be similar, since both sites are rich in sulfide, methane and DIC, and show sulfate depletion concomitant with methane accumulation. However, the distinct environmental settings and histories of these two cores, at the heavily compacted, seep-influenced base of the Sonora Margin (SeepP06), and under the oxygen minimum zone waters of the upper Sonora Margin (OMZP12), ultimately select for different archaeal communities.

## Conclusion

In the greater Guaymas Basin and Sonora Margin area, complex geological and oceanographic processes impose environmental controls on different sedimentary habitats and their archaeal populations relative to background control sites. In background sediments, archaeal communities vary little with depth after the surface/subsurface transition; here, subsurface communities result primarily from long-term survival likely conferred by relatively reduced mortality (Kirkpatrick et al., 2019). In contrast, localized factors, including water column anoxia, methane seepage, and hydrothermal circulation, constrain the biodiversity and potential biogeochemical activity of sedimentary Archaea across our benthic survey in specific ways. Local sediment biogeochemistry has to be viewed in a broader context – within the history and evolution of a particular site – to reveal its influence on selective survival for certain lineages and subsequent shaping of the resident archaeal ecosystem.

## Materials and Methods

### Sample Collection

All samples were collected using piston coring during R/V *El Puma* (Universidad Nacional Autónoma de México, UNAM) Expedition Guaymas14 to the Gulf of California, October 14-27^th^, 2014. A 5-m long piston core (RNVP11) was obtained on Oct 21, 2014 from the central basin within the ring (27°N30.5090/111°W40.6860, 1749 m; core length 4.9 m), parallel to a control core (ContP10) approx. 1 mile to the west of Ringvent (27°N30.5193/111°W42.1722; 1731 m depth, 3.93 m core length) collected on the same day. Core SeepP06 was obtained on Oct. 19 from the lower Sonora Margin, near its boundary with the Ridge flanks (27°N38.8367/111°W36.8595; 1681 m depth, 3.95 m core length). Core OMZP12 was taken on Oct. 22 from the upper Sonora Margin (27°N52.1129/111°W41.5902, 667 m, 4 m core length) in the oxygen minimum zone as previously determined by water column oxygen profiling (Calvert, 1964). Core ContP03 was collected on Oct. 17 from the northwestern end of the ridge flanks (27°N37.6759/ 111°W52.5740; 1611 m depth, 3.27 m core length. Core ContP13 was obtained on Oct. 22 from the southeastern ridge flank of Guaymas Basin (27°N12.4470/111°W13.7735, 1859m depth, 3.31 m core length).

### Geochemical Analyses

Porewater was obtained from freshly collected sediments on RV *El Puma* by centrifuging ca. 40 ml sediment samples in 50 ml conical Falcon tubes for ca. 5 to 10 minutes, using a Centra CL-2 Tabletop centrifuge (Thermo Scientific) at approx. 1000*g*, until the sediment had settled and produced ca. 8 to 10 ml of porewater. Porewater was extracted from 5 cm thick sediment samples, which are designated by the top of each sample. For example, a “95 cm” geochemistry sample extends from 95 to 100 cm below the sediment surface. Sulfate, sulfate, methane and DIC porewater profiles for cores SeepP06 to OMZP12 were previously published (Teske et al., 2019), and are re-plotted here for comparison with unpublished profiles from cores ContP03 and ContP13. Porewater analyses were performed as previously described, using the colorimetric cline assay for sulfide, ion chromatography for sulfate, and GC-IRMS for DIC and methane (Teske et al., 2019). Carbon and nitrogen isotopic and elemental composition was determined at the Stable Isotope Laboratory (SIL) at the University of California, Santa Cruz (UCSC). Bulk sediment δ^15^N and elemental ratio data were collected using 20mg samples in Sn capsules; organic δ^13^C and elemental composition data were collected using 2.5mg samples of acidified sediment in Sn capsules. All samples were measured by Dumas combustion performed on a Carlo Erba 1108 elemental analyzer coupled to a ThermoFinnigan Delt Plus XP isotope ratio mass spectrometer (EA-IRMS). An in-house gelatin standard, Acetanilide, and an in-house bulk sediment standard, “Monterey Bay Sediment Standard”, were used in all runs. Reproducibility of an in-house matrix-matched sediment standard is <0.1‰ VPDB for δ^13^C and <0.2‰ AIR for δ^15^N. Data is corrected for blank, and for drift when appropriate. Carbon and nitrogen elemental composition was estimated based on standards of known composition, for which analytical precision is determined to be better than 1 %. Filtered but unamended porewater samples, stored at 4°C, were used for quantifying multiple stable ions, including silicate, by ion chromatography at GEOMAR, Kiel, Germany (Hensen et al., 2007). All geochemical data in this study are publicly available at the Biological and Chemical Oceanography Data Management Office (BCO-DMO) under the following dataset IDs: 661750, 661658, 66175 and 661808 for Methane, DIC, Sulfate and Sulfide, respectively.

### 3. DNA extraction and gene sequencing

Samples for DNA sequencing [approx. 2 cm^3^ each] were obtained by syringe coring at the indicated depth [in cm] below the sediment surface. DNA for all survey sites was extracted from ~0.5-1.0 cm^3^ sediment sample volumes using the Powersoil DNA extraction kit according to the manufacturer’s instructions (QIAGEN, Carlsbad, CA, USA). Archaeal 16S rRNA gene amplicons from DNA extracts were generated using the following primer set: A751F: 5’-CGA CGG TGA GRG RYG AA-3’ and A1204R: 5’-TTM GGG GCA TRC NKA CCT-3’(Baker et al., 2003). Amplicons were sequenced on an Illumina MiSeq platform (Illumina, San Diego, CA, USA) at the Center for Biofilm Engineering in Bozeman, Montana. Sequencing run specifications are found in the Visualization and Analysis of Microbial Population Structures (VAMPSs) website (https://vamps.mbl.edu/resources/primers.php) (Huse et al., 2014).

### 4. Sequence Processing

Sequences were processed with *mothur* v.1.39.5 (Schloss et al., 2009) following the *mothur* Illumina MiSeq SOP (Kozich et al., 2013). Briefly, forward and reverse reads were merged into contigs and selected based on primer-specific amplicon length and the following parameters: maximum homopolymers of 6bp, and zero ambiguities. High quality sequences were aligned against the *mothur*-recreated Silva SEED v132 database (Yarza et al., 2010) and subsequently pre-clustered at 1% dissimilarity. As suggested elsewhere (Kozich et al., 2013), spurious sequences are mitigated by abundance ranking and merging with rare sequences based on minimum differences of three base pairs. Chimeras were detected and removed using UCHIME de novo mode (Edgar et al., 2011). Sequences were then clustered, by generating a distance matrix using the average neighbor method, into operational taxonomic units (OTUs, 97% similarity cutoff). OTU classification was performed on *mothur* using the SILVA v132 database as implemented using the classify.seqs command using the Wang algorithm (kmer assignment with 1/8 kmer replacement as bootstrap) and cutoff=80 (minimal bootstrap value for sequence taxonomy assignment). All sequence data are publically available at the following repository: NCBI under BioProject PRJNA553578 and accession numbers SRX6444849-SRX6444877.

### 5. Sequence Analyses

#### 5.1 Community Analyses and Visualizations

Community analyses were performed in *RStudio* version 0.98.1091 (Racine, 2012), implemented in R version 3.5.2, using the *vegan* (Oksanen et al., 2015) and *phyloseq* (McMurdie and Holmes, 2013) R-packages. Sample richness analyses used the R package *breakaway* (Willis et al. 2017) for inferring precision of diversity estimations given the heterologous sequencing depth. Data were rlog normalized using *DESeq2* (Love et al., 2014) prior to ordination using Bray-Curtis distances. An identical normalization strategy was used on Bray-Curtis distances for co-occurrence network analysis performed using the makenetwork() *phyloseq* command and visualized using the *igraph* R-package. DESeq2 was also used to perform differential abundance analyses of taxa with low abundance taxa (*n* < 100 total reads per OTU) removed for the un-rarefied dataset, as suggested elsewhere (McMurdie and Holmes, 2014).

#### 5.2 Phylogenetic Analyses

Sequence alignments were performed using the high speed multiple sequence alignment program MAFFT (Katoh and Standley, 2013) with the command: mafft --maxiterate 1000 – localpair seqs.fasta > aligned.seqs.fasta. Maximum likelihood trees with 100 bootstrap support were constructed using the RAxML (Stamatakis, 2014) program using the following parameters: raxmlHPC -f a -m GTRGAMMA -p 12345 -x 12345 -# 100 -s aligned.seqs.fasta -n T.tree, -T 4 ML search + bootstrapping. Newick trees files were uploaded to FigTree v1.4.2 for visualization.

## Data availability

Geochemical data are available at the BCO-DCO under these reference links:

Porewater methane data: https://www.bco-dmo.org/dataset/661750/data

Porewater sulfate data: https://www.bco-dmo.org/dataset/661775/data

Porewater DIC data: https://www.bco-dmo.org/dataset/661658/data

Porewater sulfide: https://www.bco-dmo.org/dataset/661808/data

## Acknowledgements

Sampling in Guaymas Basin was funded by NSF OCE grant 1449604 “Rapid Proposal: Guaymas Basin site survey cruise for IODP proposal 833” to AT; NSF C-DEBI grant “Characterizing subseafloor life and environments in Guaymas Basin” to AT and ACR. This is C-DEBI Publication No. XXX. Thanks go to Zachary Stewart for helping with the sedimentary organic matter geochemistry analyses. We thank the Science crew of R/V *El Puma* for excellent piston coring skills and a very enjoyable cruise.

## Author Contributions

APT conceived the study. LJM, ACR, and APT collected samples. CM led the *El Puma* cruise and the piston coring effort. AB extracted DNA from sediments. LJM performed sequencing in the lab of MWF. ACR, LJM and CH performed biogeochemical and sedimentological analyses. GAR analyzed the sequence data with phylogeny input from APT. GAR and APT wrote the manuscript with input from all authors.

