## Supplemental Information for "The Guaymas Basin Subseafloor Sedimentary Archaeome Reflects Complex Environmental Histories"

Author affiliations:
1: Department of Marine Sciences, University of North Carolina at Chapel Hill, NC, USA.
2: College of Veterinary Medicine, Western University of Health Sciences, Pomona, CA, USA.
3: Department of Land Resources and Environmental Sciences, Montana State University, Bozeman, MT, USA.
4: Center for Biofilm Engineering, Montana State University, Bozeman, MT, USA.
5: Department of Microbiology and Immunology, Montana State University, Bozeman, MT, USA.
6: Instituto de Geofisica, Universidad Nacional Autónoma de México, Coyoacán, Mexico.
7: GEOMAR Helmholtz Centre for Ocean Research, Kiel, Germany.
8: Ocean Sciences Department, University of California, Santa Cruz, CA, USA.

*: Corresponding author

Correspondence:
Dr. Gustavo A. Ramírez




| ***Core ID*** | **Latitude** | **Longitude** | **Collection Date (2014)** | **Core Length (m)** | **Water Depth (m)** |
|---|---|---|---|---|---|
| ***ContP3*** | 27°N37.6759 | 111°W52.5740 | Oct. 17 | 3.27 | 1611 |
| ***SeepP6*** | 27°N38.8367 | 111°W36.8595 | Oct. 19 | 3.95 | 1681 |
| ***ContP10*** | 27°N30.5193 | 111°W42.1722 | Oct. 21 | 3.93 | 1731 |
| ***RNVP11*** | 27°N30.5090 | 111°W40.6860 | Oct. 21 | 4.9 | 1749 |
| ***OMZP12*** | 27°N52.1129 | 111°W41.5902 | Oct. 22 | 4 | 667 |
| ***ContP13*** | 27°N12.4470 | 111°W13.7735 | Oct. 22 | 3.31 | 1859 |

**Table S1.** Core site metadata.

| Sample Name | DNA yield (ng/µL) | Num. of seqs post Mothur QC and chimera removal |
|---|---|---|
| ContP3_9 | 7 | 21,443 |
| ContP3_104 | 6.9 | 47,239 |
| ContP3_202 | 6.6 | 16,038 |
| ContP3_301 | 9.4 | 45,559 |
| SeepP6_5 | 9 | 17,196 |
| SeepP6_105 | 4.3 | 11,595 |
| SeepP6_205 | 9.1 | 9,274 |
| SeepP6_304 | 9.4 | 18,043 |
| SeepP6_394 | 8 | 10,047 |
| ContP10_5 | 9.2 | 25,975 |
| ContP10_104 | 7.7 | 12,289 |
| ContP10_204 | 8 | 35,076 |
| ContP10_303 | 14.5 | 29,782 |
| ContP10_378 | 7.6 | 25,682 |
| RNVP11_5 | 6.7 | 11,184 |
| RNVP11_95 | 6.7 | 30,452 |
| RNVP11_195 | 7.1 | 2,978 |
| RNVP11_295 | 7 | 19,515 |
| RNVP11_394 | 7.4 | 14,142 |
| RNVP11_468 | 7.9 | 29,851 |
| OMZP12_5 | 7.9 | 63,690 |
| OMZP12_105 | 9 | 51,384 |
| OMZP12_204 | 7.8 | 167,234 |
| OMZP12_304 | 7.3 | 154,763 |
| OMZP12_379 | 8.1 | 76,729 |
| ContP13_5 | 6.6 | 17,573 |
| ContP13_111 | 7.9 | 47,432 |
| ContP13_210 | 6.8 | 25,989 |
| ContP13_310 | 7.3 | 24,873 |

**Table S2**. Total DNA yield and high quality sequence numbers for all samples.

| Core_cmbsf | All_ANME | ANME-1 | ANME-2a-2b | ANME-2c |
|---|---|---|---|---|
| ContP3_005 | 0.034 | 0.000 | 0.000 | 0.034 |
| ContP3_104 | 0.002 | 0.002 | 0.000 | 0.000 |
| ContP3_202 | 0.000 | 0.000 | 0.000 | 0.000 |
| ContP3_301 | 0.000 | 0.000 | 0.000 | 0.000 |
| SeepP6_005 | 0.030 | 0.018 | 0.012 | 0.000 |
| SeepP6_105 | 8.863 | 8.863 | 0.000 | 0.000 |
| SeepP6_205 | 32.063 | 32.063 | 0.000 | 0.000 |
| SeepP6_304 | 32.446 | 32.440 | 0.006 | 0.000 |
| SeepP6_394 | 39.810 | 39.810 | 0.000 | 0.000 |
| ContP10_005 | 0.111 | 0.088 | 0.024 | 0.000 |
| ContP10_104 | 0.092 | 0.092 | 0.000 | 0.000 |
| ContP10_204 | 0.003 | 0.000 | 0.003 | 0.000 |
| ContP10_303 | 0.447 | 0.447 | 0.000 | 0.000 |
| ContP10_378 | 0.000 | 0.000 | 0.000 | 0.000 |
| RNVP11_005 | 0.009 | 0.009 | 0.000 | 0.000 |
| RNVP11_095 | 0.000 | 0.000 | 0.000 | 0.000 |
| RNVP11_195 | 0.988 | 0.988 | 0.000 | 0.000 |
| RNVP11_295 | 0.000 | 0.000 | 0.000 | 0.000 |
| RNVP11_394 | 0.000 | 0.000 | 0.000 | 0.000 |
| RNVP11_486 | 0.000 | 0.000 | 0.000 | 0.000 |
| OMZP12_005 | 0.000 | 0.000 | 0.000 | 0.000 |
| OMZP12_105 | 0.123 | 0.121 | 0.002 | 0.000 |
| OMZP12_204 | 2.098 | 2.098 | 0.000 | 0.000 |
| OMZP12_304 | 0.629 | 0.629 | 0.000 | 0.000 |
| OMZP12_378 | 0.967 | 0.967 | 0.000 | 0.000 |
| ContP13_005 | 0.476 | 0.429 | 0.029 | 0.018 |
| ContP13_111 | 0.006 | 0.002 | 0.004 | 0.000 |
| ContP13_211 | 0.055 | 0.012 | 0.043 | 0.000 |
| ContP13_311 | 0.004 | 0.000 | 0.004 | 0.000 |

**Table S3.** Percent of total community contribution of ANME sequences in all samples based on SILVA132 taxonomic assignments. The All_ANME column shows the percent contribution of sequences classified as ANME in each sample. Columns ANME-1, ANME-2a-2b, and ANME-2c show the percent breakdown of the respective ANME lineages in each sample and their sum is equal to the All_ANME column percentage.

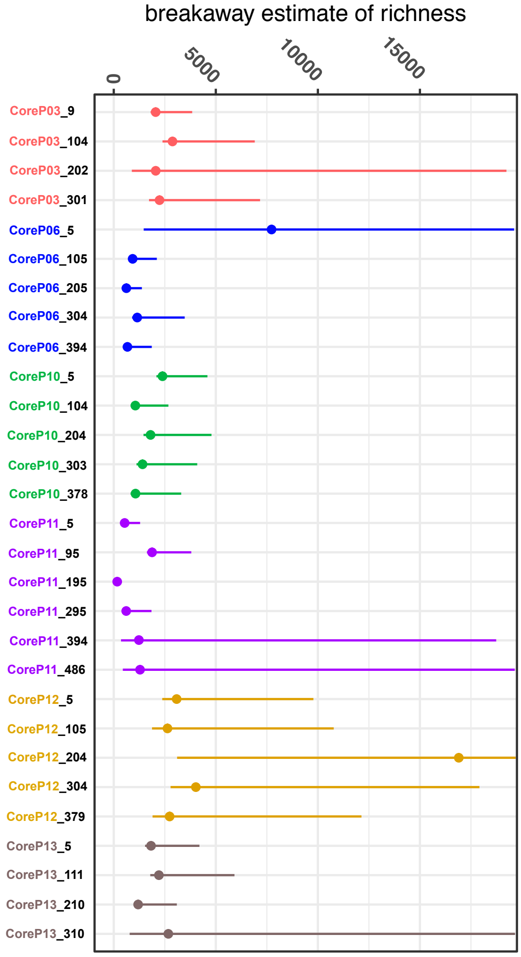


**Figure S1**. Breakaway() estimate of total species richness with model confidence intervals for color-coded cored site for all depths.

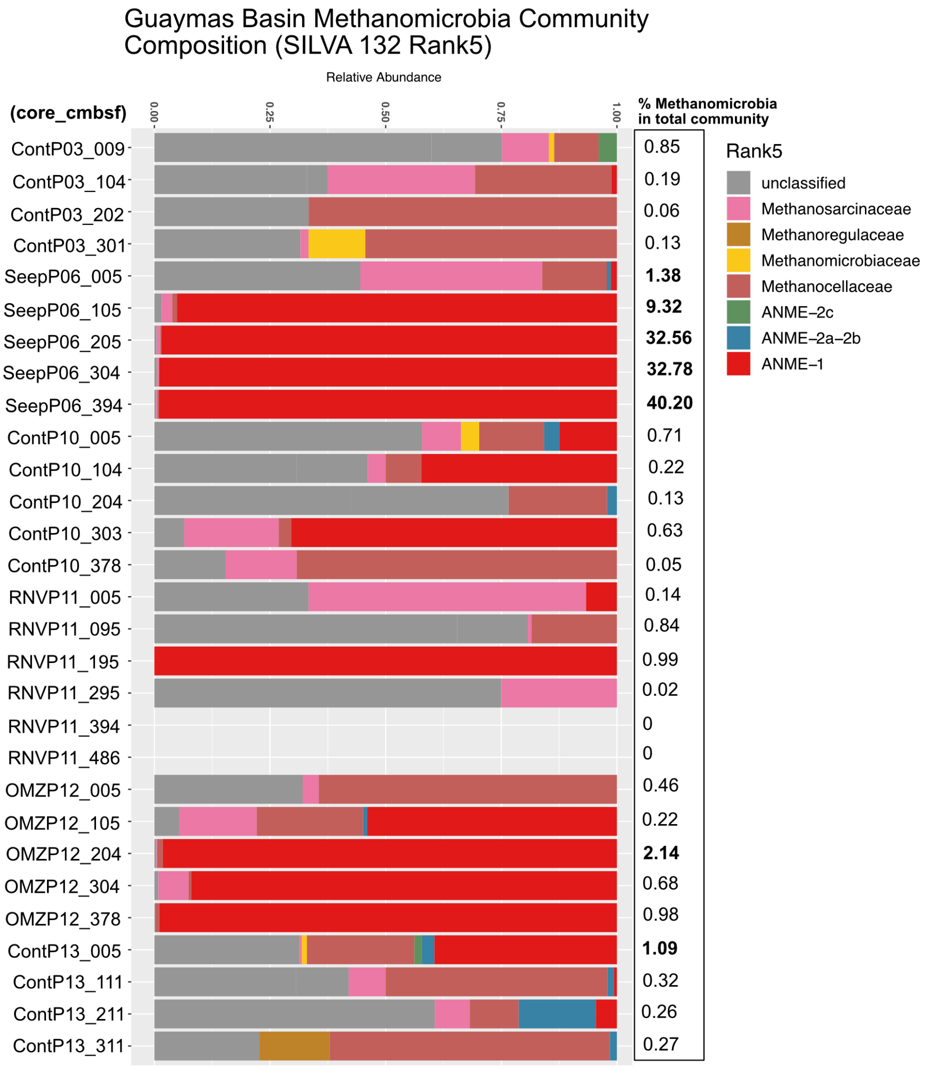


**Figure S2.** Methanomicrobia community composition for all cores in this survey.
